# Locating and Quantifying Carbon Steel Corrosion Rates Linked to Fungal B20 Biodiesel Degradation

**DOI:** 10.1101/2021.06.29.450458

**Authors:** James G. Floyd, Blake W. Stamps, Wendy J. Goodson, Bradley S. Stevenson

## Abstract

Fungi that degrade B20 biodiesel in storage tanks have also been linked to microbiologically influenced corrosion (MIC). A member of the filamentous fungal genus *Byssochlamys*, and a yeast from the genus *Wickerhamomyces* were isolated from heavily contaminated B20 storage tanks from multiple Air Force bases. Although these taxa were linked to microbiologically influenced corrosion *in situ*, precise measurement of their corrosion rates and pitting severity on carbon steel was not available. In the experiments described here, we directly link fungal growth on B20 biodiesel to higher corrosion rates and pitting corrosion of carbon steel under controlled conditions. When these fungi were growing solely on B20 biodiesel for carbon and energy, consumption of FAME and n-alkanes was observed. The corrosion rates for both fungi were highest at the interface between the B20 biodiesel and the aqueous medium, where they acidified the medium and produced deeper pits than abiotic controls. *Byssochlamys* produced the most corrosion of carbon steel and produced the greatest pitting damage. This study characterizes and quantifies the corrosion of carbon steel by fungi that are common in fouled B20 biodiesel through their metabolism of the fuel, providing valuable insight for assessing MIC associated with storage and dispensing B20 biodiesel.

**IMPORTANCE:** Biodiesel is widely used across the United States and worldwide, blended with ultralow sulfur diesel in various concentrations. In this study we were able to demonstrate that the filamentous fungi *Byssochlamys* AF004 and the yeast *Wickerhamomyces* SE3 were able to degrade fatty acid methyl esters and alkanes in biodiesel causing increases in acidity. Both fungi also accelerated the corrosion of carbon steel, especially at the interface of the fuel and water, where their biofilms were located. This research provides controlled, quantified measurements and the localization of microbiologically influenced corrosion caused by common fungal contaminants in biodiesel fuels.

## INTRODUCTION

Biodiesel production within the United States (U.S.) greatly expanded in response to high petroleum prices and an increased need for energy security after the September 11, 2001 terrorist attacks (1, 2). Over two decades afterwards, world biodiesel consumption has continued to increase, reaching 9.3 billion gallons in 56 countries in 2016 (3). Consumption in the U.S. increased from 322 million gallons annually in 2009 to 1.8 billion gallons in 2019 (4). The U.S. increase in biodiesel consumption was largely driven by an increased interest in energy security but there are other advantages to its’ use. When biodiesel is blended with or used as an additive to ultra-low sulfur diesel (ULSD), fuel lubricity is restored and emissions of carbon and particulates are partially offset (5, 6). The increased energy independence and fuel performance resulted in widespread adoption of biodiesel blends worldwide. The largest energy consumer in the U.S., the United States Department of Defense (DoD), sought to diversify its’ energy supplies to reduce fuel security risk through the use of renewable ‘drop-in’ fuels to replace unblended fuels such as ULSD (7). The United States Air Force (USAF), a service within the DoD, rapidly implemented the use of a drop-in fuel in the form of a 20% first-generation biodiesel blend (B20) in non-tactical ground vehicles and equipment. B20 is composed of a 20:80 volume/volume blend of fatty acid methyl esters (*i*.*e*. FAME biodiesel) and ULSD, which is compatible with existing engines and storage infrastructure (8). The FAME used as biodiesel is a renewable resource produced through an esterification reaction with animal and plant triglycerides and methanol (9). However, the advantages that adoption of B20 represent are tempered by potential limitations.

Biodiesel is more hygroscopic and oxidatively unstable compared to petroleum-based diesel, making fuels containing biodiesel more susceptible to microbiological degradation (10). This is especially true during long term storage as microorganisms can readily degrade the FAME in B20 (11). The half-life of the FAME in B20, as well as C7 to C20 alkanes were 2.1 to 2.8 days in aqueous microcosm experiments, suggesting that both the FAME and alkanes in biodiesel blends is a viable oxidizable substrate for microbial growth (12). The biodegradation of FAME is carried out through the β- oxidation pathway, which involves the sequential removal of two-carbon components and production of acetic acid. Acetic acid can then be converted to acetyl-CoA and used in the Krebs cycle or exported from the microbial cell (13–15). Methanol produced from the de-esterification of FAME is readily metabolized by acid-tolerant fungi and bacteria (16). The aliphatic alkanes in B20 are also readily degraded by both fungi and bacteria suggesting that the oxidation of B20 under aerobic conditions is highly favorable (17, 18). Under aerobic conditions alkanes are oxidized by alkane monooxygenases producing a fatty acid that is shuttled into the β-oxidation pathway, producing additional acetic acid (19). Free fatty acids and acidic byproducts produced from the metabolism of many of the components of B20 fuel can acidify both the fuel and any water present within the storage tank, inducing or accelerating corrosion, leading to damaged vehicles and storage infrastructure.

Corrosion directly or indirectly caused by microorganisms is known as microbiologically influenced corrosion, or MIC. Annually, corrosion costs an estimated 2.5 trillion dollars worldwide, up to 20% of which is attributed to MIC (20). MIC is initiated at the interface of microorganisms, often in the form of a biofilm, and material surfaces. Biofilms affect the physical and chemical environment of metallic surfaces, impacting the kinetics of cathodic or anodic reactions (21, 22). Microorganisms in biofilms can also secrete enzymes that attack metals, increase local acidity, create differential aeration, and form galvanic cells which accelerate corrosion under otherwise aerobic conditions (23).

Fungi can increase corrosion rates of mild steel when grown on ULSD as a sole carbon and energy source (24). Fungi grow more rapidly and produce more biomass on the FAME in biodiesel blends than ULSD, which could lead to a greater corrosion risk to infrastructure storing or dispensing fuels containing biodiesel (25). Filamentous fungi such as *Penicillium* and *Aspergillus* increase the rate of steel corrosion when degrading diesel fuel (26). It was speculated that the increased steel corrosion rates from these fungi degrading diesel were likely attributed to increased oxygen concentrations in the medium caused by degrading benzene rings and aliphatic hydrocarbons to (-O-CH_2_-). The authors also acknowledged that these organisms were likely producing organic acids that could also play a role in the increased corrosion risk. Additionally, studies have shown that fungi contaminating biodiesel storage tanks in Brazil were able to degrade fuels containing 5%, 10%, 20% and 100% biodiesel (25). Although they did not measure corrosion during their investigation, the authors did note that as fungi degraded these fuels, there was an increase in acidity that could impact corrosion risks.

There is a critical need to provide quantified rates and measurements of how fungi can degrade fuels, such as B20, and subsequently contribute to MIC in critical fuel storage infrastructure.

When the USAF started using B20 to meet mandated energy requirements, problems quickly arose (27). Numerous USAF bases reported particulates in the fuel from B20 storage tanks and fouled filters on dispensers (28). Subsequent *in-situ* analysis of corrosion confirmed that tanks with obvious fungal contamination had significant pitting corrosion relative to controls. These storage tanks being monitored were still in operation making it difficult to control for other factors that might have contributed to the perceived increase in corrosion risk. Herein we describe the isolation of organisms responsible for fouling contaminated B20 storage tanks and directly link them to fuel degradation and increased corrosion risks. To address our lack of a direct causative link between the previously identified fungal taxa within USAF B20 tanks and their ability to both increase corrosion rates of carbon steel and degrade B20, members of the abundant taxa *Byssochlamys* and *Wickerhamomyces*, were isolated. These isolates were used to determine their ability to metabolize B20 biodiesel and spatially induce corrosion of carbon steel.

## Results

### Characterization and Phylogenetic Identification of Fungi Isolated from B20 Biodiesel

The fungal ITS sequences that were sequenced suggested that the filamentous fungal isolate was most closely related to *Byssochlamys nivea* (Fig. 1A) and the yeast isolate was most closely related to *Wickerhamomyces anomalus* (Fig. 1B).

**FIG 1:**
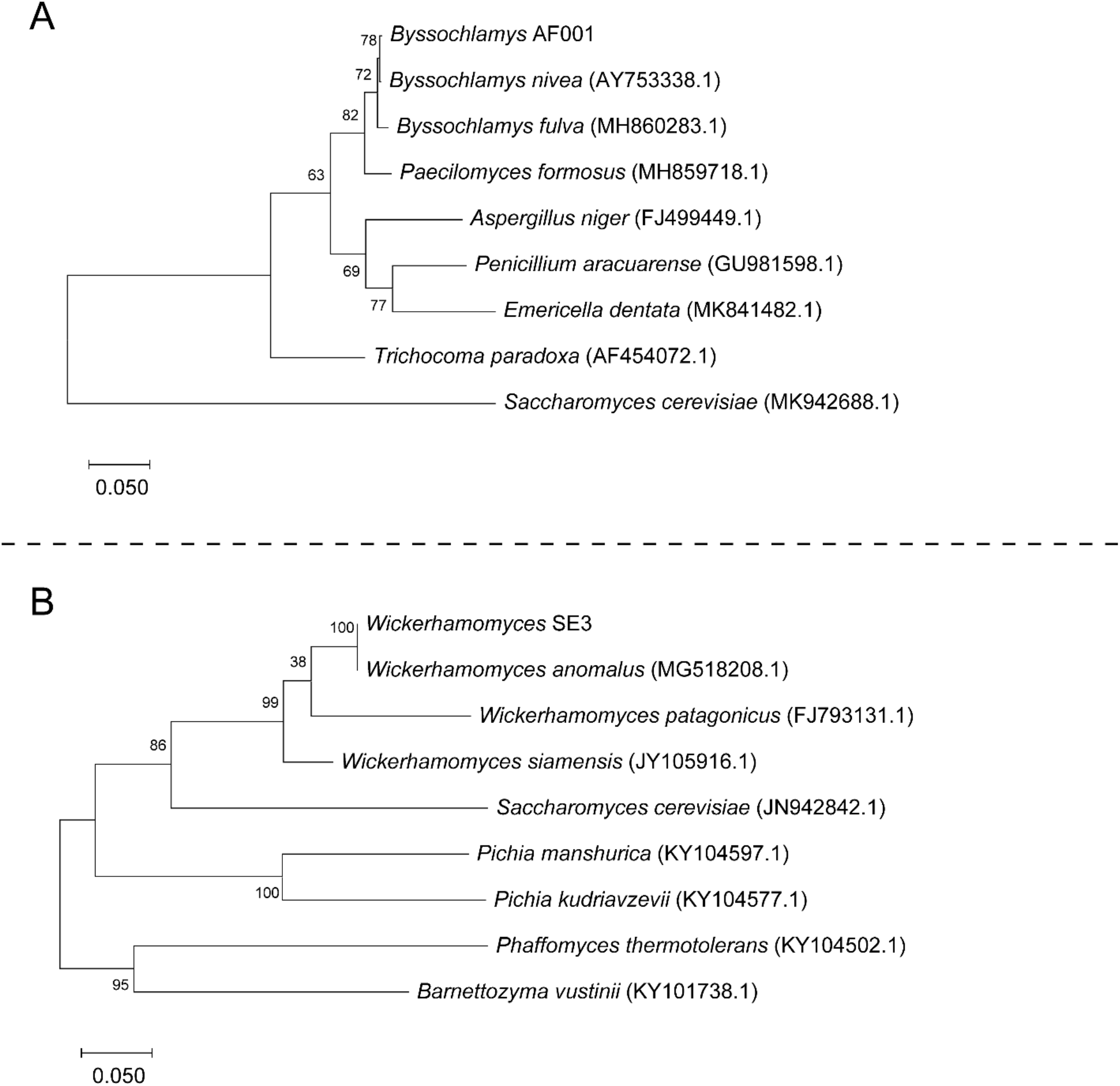
Maximum likelihood tree based on ITS sequence similarity among close phylogenetic relatives (NCBI accession numbers in parentheses) of the *Byssochlamys* sp. AF001 (**A**) and *Wickerhamomyces* SE3 (**B)** isolates. Bootstrap values above 50 percent for 500 samples are shown at relevant nodes.

### Fungal Growth in Bioreactors

*Byssochlamys* AF001 and *Wickerhamomyces* SE3 were presumably in lag phase and had no apparent growth during the first 7 days in the bioreactors. However, after 14 days the MPN of *Byssochlamys* on the surface of the carbon steel witness coupons had increased over an order of magnitude, the density of *Wickerhamomyces* on the coupon surfaces and the liquid medium had increased in a similar fashion based on CFUs (Fig 2).

**FIG 2.**
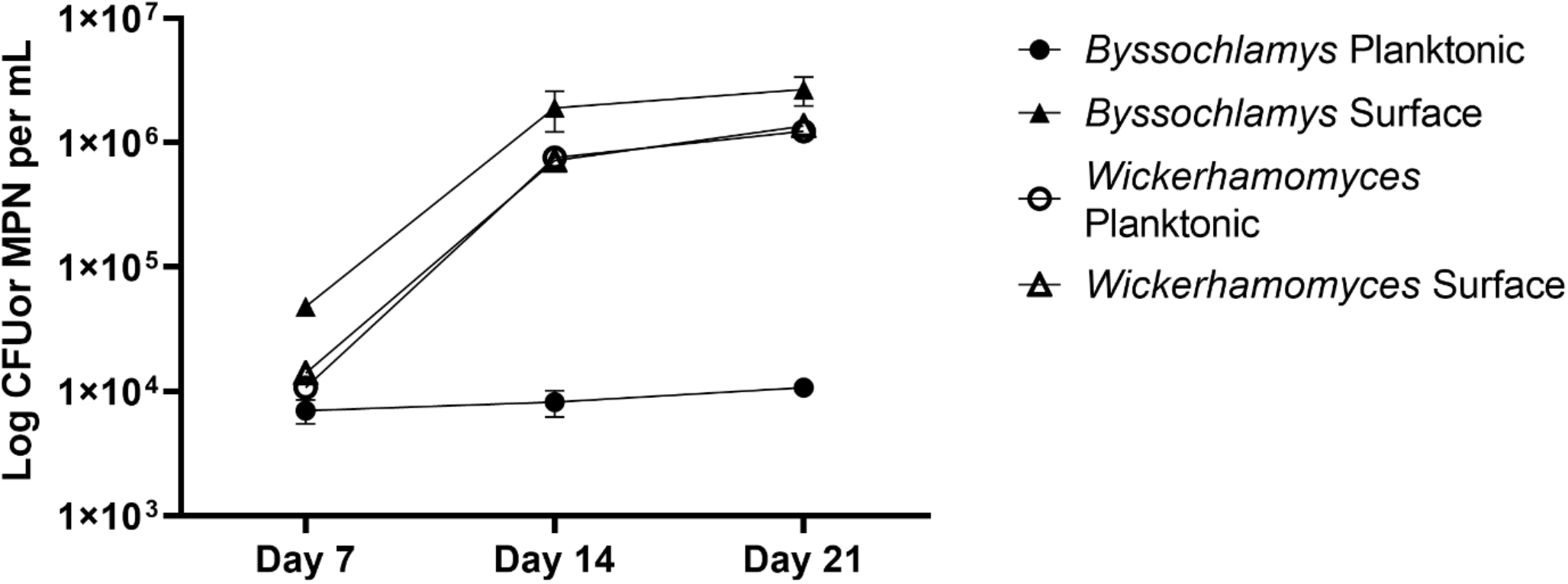
Density of planktonic populations (circles) and biofilms (squares) in bioreactors inoculated with *Byssochlamys* AF001 (MPNs; Black) or *Wickerhamomyces* SE3 (CFUs; Gray). Error bars represent 95% confidence intervals for mean *Byssochlamys* AF001 MPNs or *Wickerhamomyces* SE3 CFUs (n=3).

**FIG 3.**
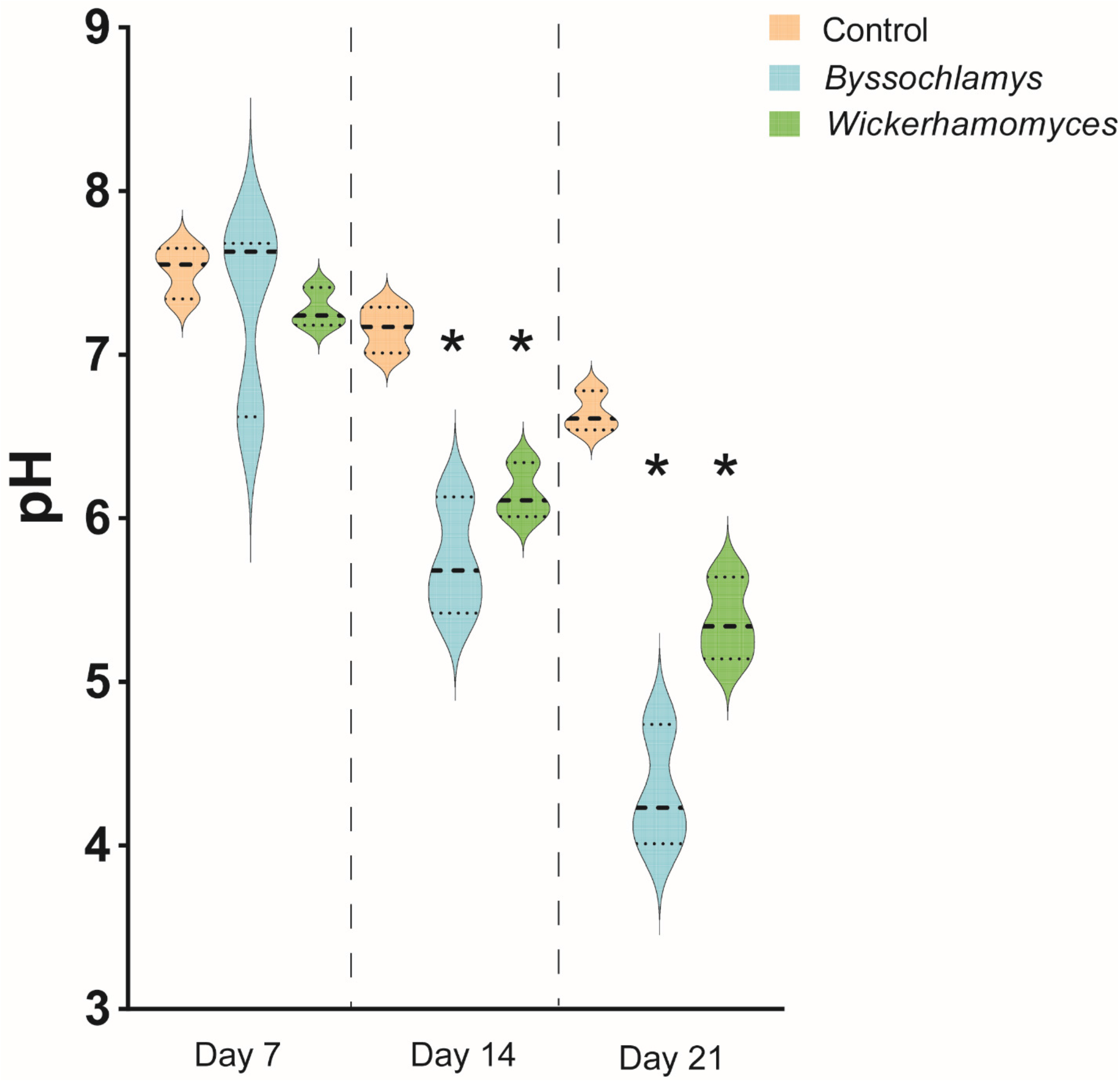
pH values of the aqueous phase in bioreactors inoculated with *Byssochlamys* AF001(Blue), *Wickerhamomyces* (Green), or controls (Tan) over time. Bold dashed lines represent the median while the nonbold dashed lines represent the data quartiles. Asterisks indicate a significant difference between the pH of inoculated and uninoculated controls.

### Acidification of Aqueous Medium

When *Byssochlamys* sp. AF001 and *Wickerhamomyces* sp. SE3 were grown on B20 biodiesel as the sole carbon and energy source, the pH of the aqueous phase did not decrease significantly after 7 days relative to abiotic controls (Fig. 4C). By day 14 and 21 both fungal isolates significantly reduced (p <0.05) the pH of the medium by two or more orders of magnitude compared to the abiotic controls at those times. The *Byssochlamys* isolate was responsible for the greatest reduction in pH (4.33 ± 0.31) of the aqueous phase after 21 days. The *Wickerhamomyces* isolate decreased the aqueous pH to a mean of 5.37 ± 0.21 after 21 days.

**FIG 4.**
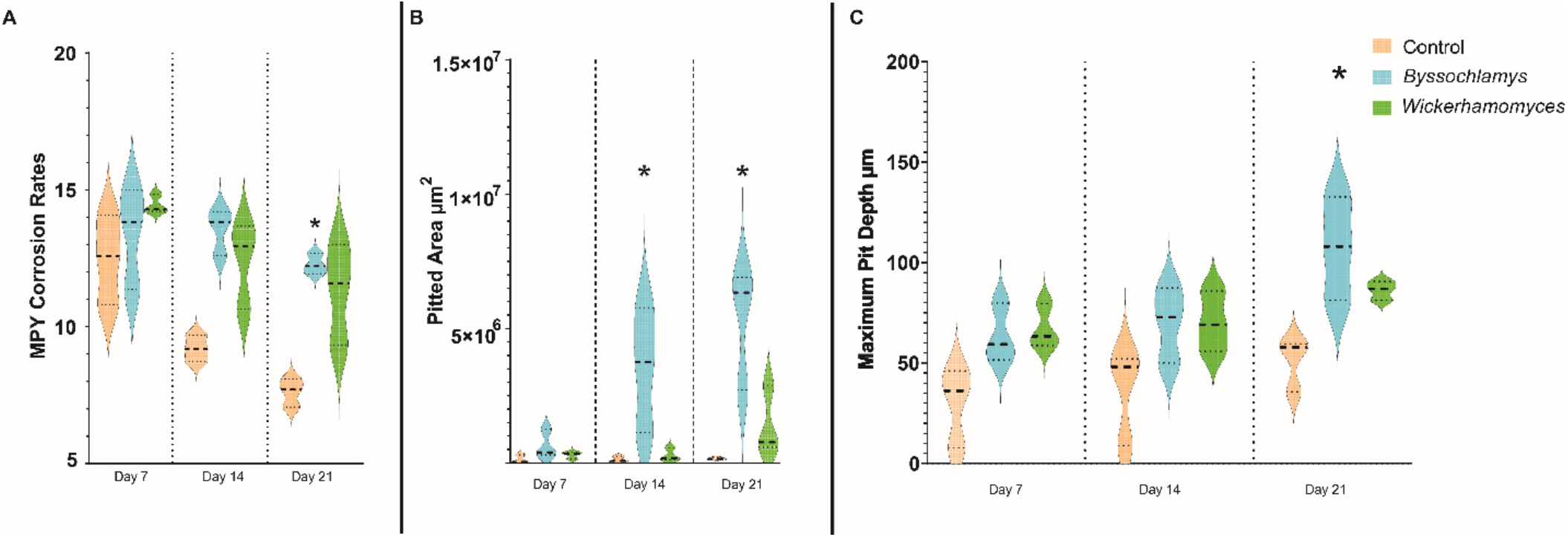
**A**. Corrosion rates in milliinches per year (MPY) of carbon steel coupons in bioreactors inoculated with *Byssochlamys* (Blue), *Wickerhamomyces* (Green), and uninoculated controls (Tan) over time. The bold dashed line represents the median (n=3) and nonbold dashed lines represent the data quartiles. Asterisks indicate significant difference between the inoculated and abiotic conditions for each time point. **B**. Total pitted area on carbon steel coupons from bioreactors inoculated with *Byssochlamys* (Blue), *Wickerhamomyces* (Green), and uninoculated controls (Tan). The bold dashed line represents the median (n=3) and nonbold dashed lines represent the data quartiles. Asterisks indicate significant difference between the inoculated and abiotic conditions for each time point. **C**. Maximum pit depths on carbon steel coupons in uninoculated bioreactors (Tan) and bioreactors inoculated with *Byssochlamys* (Blue) and *Wickerhamomyces* (Green) over time.

### Direct Measurement of Corrosion

Corrosion rates (MPY) of the carbon steel coupons by each isolate were not significantly different than abiotic controls after 7 and 14 days (Fig 4A). After 21 days, incubations with the *Byssochlamys* isolate had significantly greater (p < 0.05) corrosion rates compared to abiotic controls. *Wickerhamomyces* SE3 did not produce significantly higher corrosion rates when compared to abiotic controls. The total area pitted of witness coupons was significantly higher than that of abiotic controls only in bioreactors inoculated with *Byssochlamys* after 14 and 21 days (p <0.05) (Fig 4B). The greatest pit depth recorded for all *Byssochlamys* cultures was 132.7 µm whereas the maximum pit depth recorded for all *Wickerhamomyces* SE3 reactors was 90.7 µm.

### Colocation of biology and corrosion

In static cultures the pH of the aqueous phase was significantly lower (p <0.05) for both *Byssochlamys* and *Wickerhamomyces* compared to abiotic controls (Fig. 5A). Corrosion rates were lowest in the organic (fuel) phase and elevated in both the aqueous phase (ASW) and at the interface of fuel and medium (Fig. 6B). Corrosion rates in the fuel phase were significantly higher than abiotic controls (p<0.05), six times greater in cultures inoculated with *Byssochlamys* relative to uninoculated controls. In cultures inoculated with *Wickerhamomyces*, corrosion rates were not significantly different (p > 0.05) from controls in the fuel phase. Corrosion rates at the fuel/medium interface and in the growth medium were significantly higher for both *Byssochlamys* and *Wickerhamomyces* relative to abiotic controls (p <0.05). The greatest corrosion rates (2.1 ± 0.23 MPY *Byssochlamys* and 1.8 ± 0.36 MPY *Wickerhamomyces*) were observed at the fuel/medium interface. *Byssochlamys* produced more and deeper pitting corrosion at the fuel/medium interface compared to the uninoculated control (Fig. 7).

**FIG 5.**
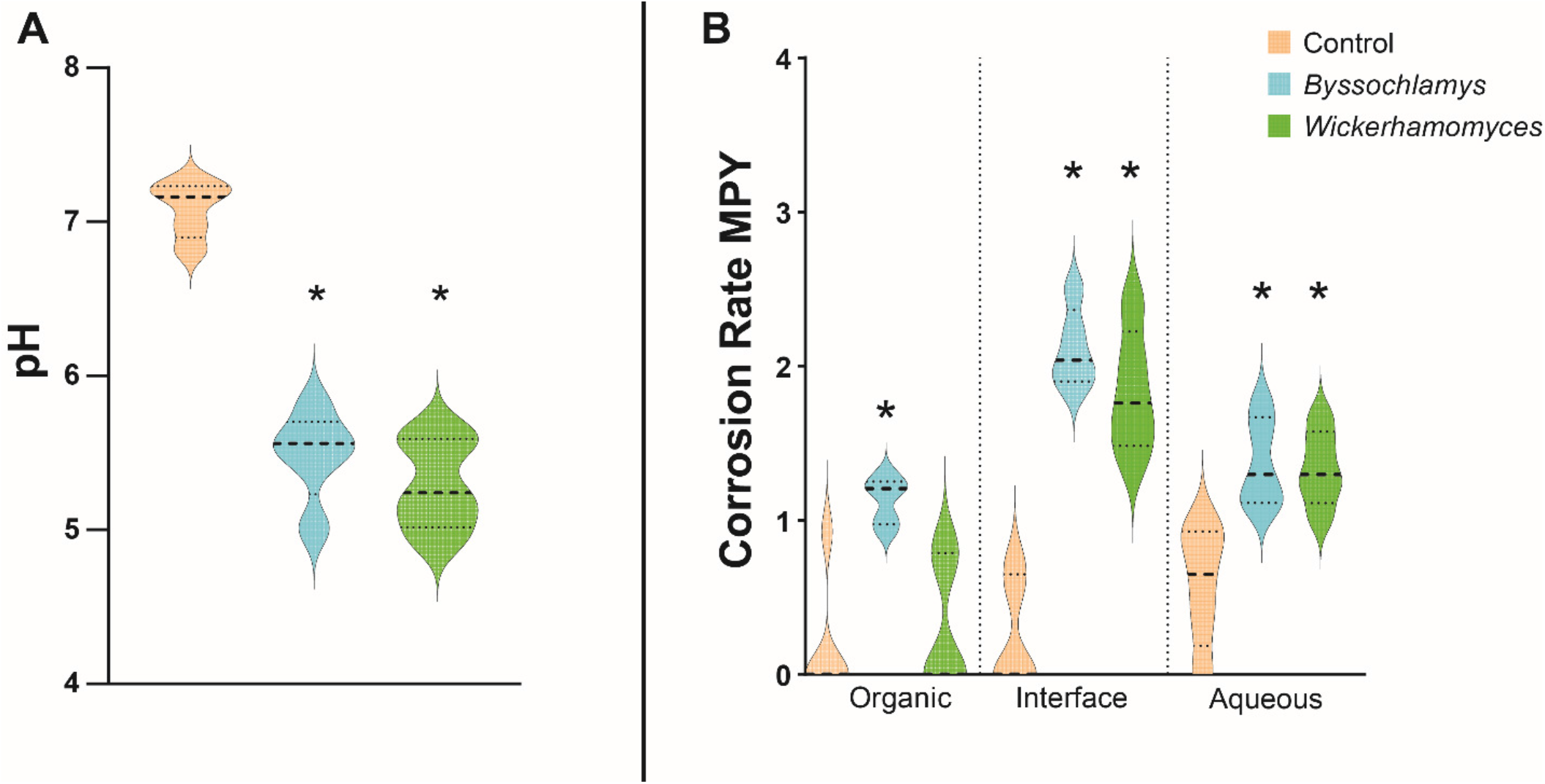
**A**. pH values of the aqueous phases after incubation of carbon steel brads with *Byssochlamys* (Blue), *Wickerhamomyces* (Green) and uninoculated controls (Tan). The bold lines represent the median and the nonbold lines on the plots represent the data quartiles (n=5). Asterisks indicate a significant difference between the inoculated and uninoculated conditions. **B**. Corrosion rates of carbon steel brads in the organic phases, organic-aqueous interface, and aqueous phase after exposure to the fungal isolates. The bold lines represent the median and the nonbold lines on the plots represent the data quartiles (n=5). Asterisks indicate a significant difference between the inoculated and uninoculated conditions.

**FIG 6.**
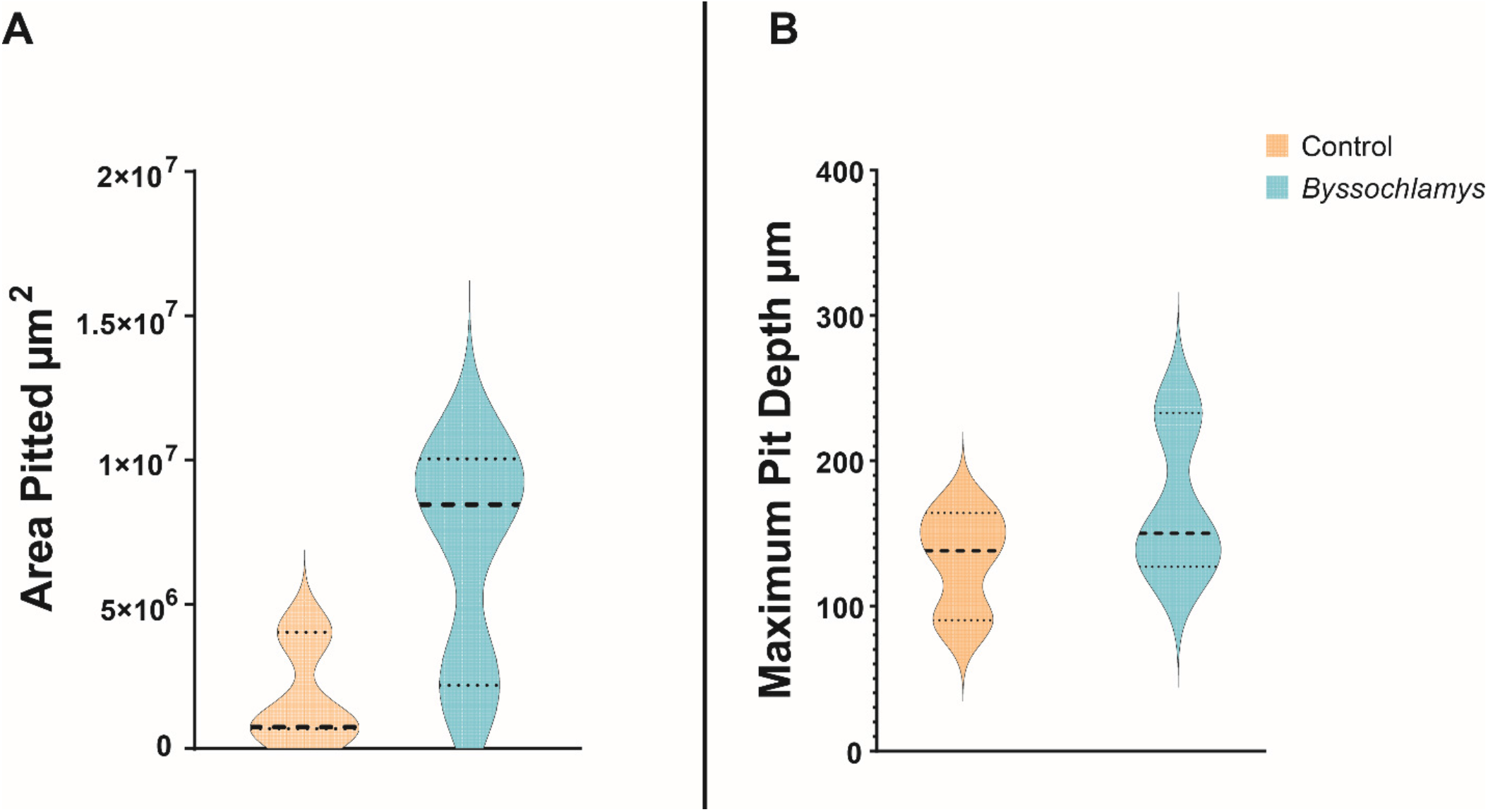
**A**. Total pitted area on carbon steel coupons (pits > 20µm below mean surface average) inoculated with *Byssochlamys* (Blue) compared to uninoculated controls (Tan). The bold line represents the medium of the data and the nonbold lines represent the data quartiles (n=3). **B**. Maximum pit depths on carbon steel coupons in abiotic controls (Tan) and flasks inoculated with *Byssochlamys* (Blue) over 90 days. The bold line represents the median of the data and the nonbold lines represent the data quartiles (n=3).

**FIG 7.**
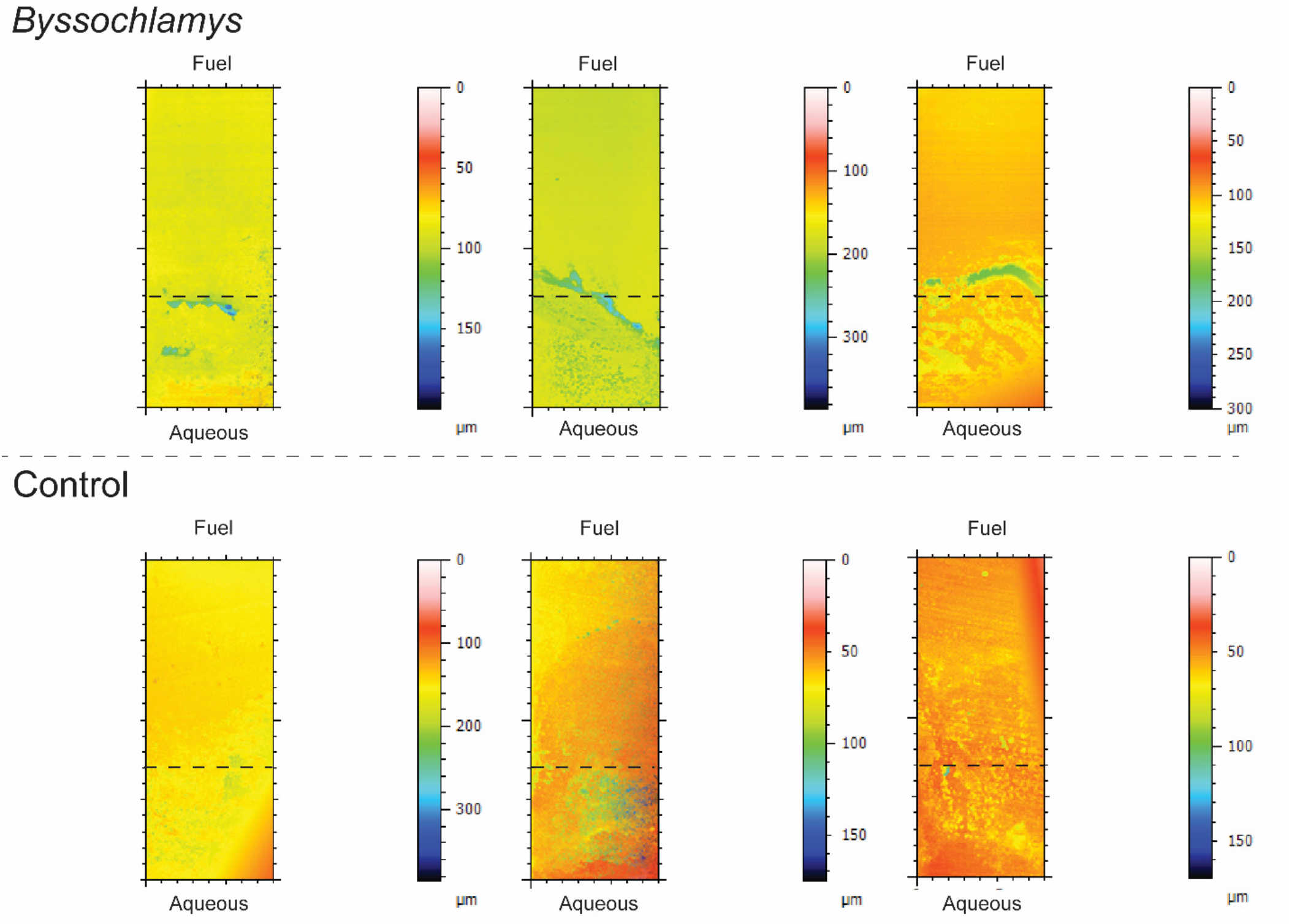
Surface depth profiles of carbon steel coupons exposed to the filamentous fungus *Byssochlamys* and uninoculated controls after 90 days. Biological replicates are show next to each other and depth was calculated from the highest point on the surface of the coupon representing 0µm. The color scale bar represents depths in microns. The dashed bar represents where the interface was located on the coupons with the fuel phase above and the aqeous phase below.

### Fuel Degradation

Both *Byssochlamys* and *Wickerhamomyces* were capable of growth in B20, metabolizing FAME and alkane components of the fuel. As a result, the peak area of all detectable FAME components and alkanes decreased over time in *Byssochlamys* and *Wickerhamomyces* cultures (Figure 8). Compared to uninoculated controls, both *Byssochlamys* and *Wickerhamomyces* cultures metabolized greater than 50 % of the cis-9-oleic acid methyl ester by day 21. Both fungi decreased the concentration of Cis-9-Oleic acid methyl ester more than any other fuel component after 21 days (measured as peak area). Analysis of the uninoculated controls over the 21 days showed evaporation of fuel components with approximately 15% reduction in the components compared to the unexposed fuel.

**FIG 8.**
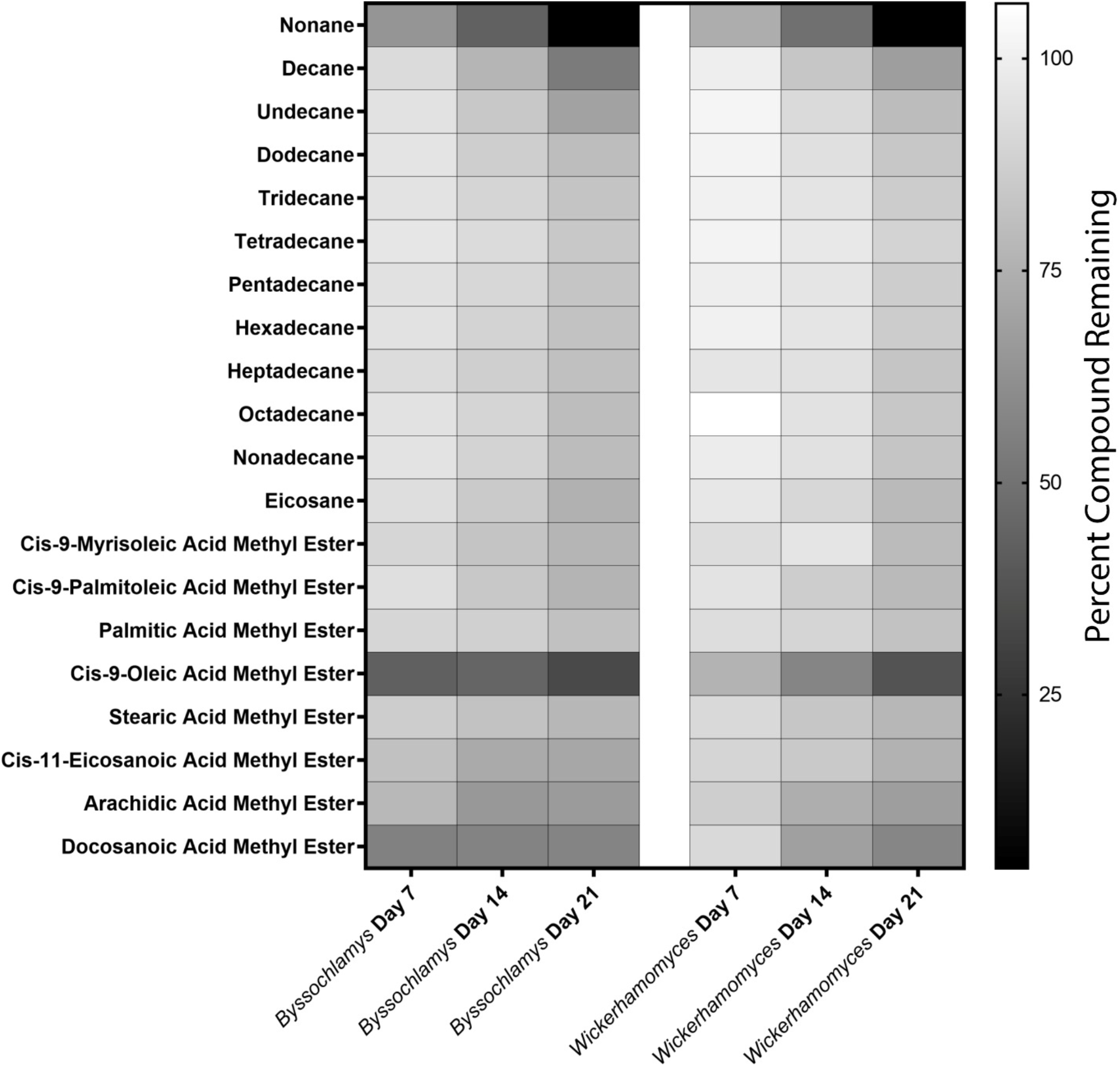
Degradation of alkanes and FAME in B20 biodiesel by *Wickerhamomyces* and *Byssochlamys* after 7, 14 and 21 days of incubation. The scale bar represents the percent of remaining compound compared to the unexposed control with white indicating no degradation and black indicating complete degradation.

## Discussion

Complex interactions between microorganisms, fuels, and metallic surfaces *in situ* make it challenging to directly link the microbial degradation of biodiesel to corrosion (29). Isolation of the relevant microorganisms and controlled laboratory experimentation is critical to linking microbial growth and metabolic activity to the corrosion of infrastructure. Both fungi isolated and identified in this study, *Byssochlamys* AF004 and *Wickerhamomyces* SE3, represented some of the most abundant fungi identified in a longitudinal study of microbial contamination and microbiologically influenced corrosion in B20 storage tanks (28). We measured the ability of these organisms to both degrade B20 and accelerate corrosion of carbon steel in controlled laboratory experiments. The co-location of microbial biomass and the greatest amount of corrosion at the interface of fuel and water provided a clear link of where the potential for corrosion would be highest in storage tanks, enhancing our ability to more specifically target regions of fuel tanks for mitigation and increasing the chance of early detection of MIC.

Two fungi that were abundant in contaminated B20 fuel samples included a filamentous fungus, *Byssochlamys*, and a yeast, *Wickerhamomyces*. Both were capable of growth on B20 as a sole carbon and energy source by degrading the readily oxidizable FAME components of B20 and several of the alkanes within 21 days. Currently the U.S. DOE and the European Union authorizes the use of B5 and B7 respectively to be added to diesel fuels (30, 31). Our work suggests that even in fuels with low concentrations of FAME such as B5 or B7 (supposedly ‘neat’ ULSD) biofouling, microbiologically influenced corrosion, and fuel degradation can become more common as biodiesel is added to petroleum diesel.

Biodegradation of both hydrocarbons and FAME in biodiesel blends produces organic acids, which acidifies both the fuel and any aqueous environments, increasing the corrosion of carbon steel (32, 33). Both fungal isolates in this study could degrade alkanes and FAME in B20 biodiesel and appeared to degrade cis-9-oleic acid methyl ester and cis-11-eicosonic acid methyl ester more readily than the other compounds. Some studies have demonstrated that both fungi and bacteria can metabolize and incorporate unsaturated fatty acids much more rapidly than saturated fatty acids and has been attributed to linear fatty acids being less efficient in entering inside cells for metabolic processes (35, 36).

Biofilms can accelerate corrosion rates by creating localized, concentrated acidic conditions at metallic surfaces or by partitioning oxygen, generating oxygen corrosion cells. Both of these mechanisms increase “pitting” or localized corrosion (34). Surface analysis conducted by white light profilometry showed that pitting corrosion was more severe on coupons exposed to the filamentous fungus *Byssochlamys* AF001 compared to the yeast *Wickerhamomyces*. This difference could be due to the thick biofilms formed by the filamentous fungus. Other filamentous fungi such as *Aspergillus niger* form thick biofilms over steel surfaces, resulting in more pitting corrosion (37). After 21 days, *Byssochlamys* biofilms generated approximately 30-fold more pitted area than the abiotic controls, and 4 times more than on coupons exposed to the yeast *Wickerhamomyces*. The elevated corrosion observed in the *Byssochlamys* cultures confirms previous studies in USAF B20 storage tanks that suggested filamentous fungi contributed to MIC (28).

While we confirmed that both fungi were capable of MIC, materials did exhibit some surface passivation, or reduction in corrosion rates over time. Corrosion rates decreased slightly over the course of the 21-day bioreactor experiment, possibly due to the formation of passivating iron oxides on the surface of the metal. This passivation could prevent further iron oxidation from occurring as rapidly as when neutral iron is exposed to the medium (38, 39). Corrosion rates plateaued over time similar to the corrosion dynamics that were observed *in situ* where *Byssochlamys* was the most abundant fungal population (40). Although overall corrosion rates decelerated over the course of 12 months, significantly more deep pits were detected, suggesting that long-term corrosion modeling remains challenging.

Both isolated fungi are capable of growth under oxic conditions, and most of the observable fungal and bacterial taxa identified *in situ* were either facultative anaerobes or known aerobic microorganisms (28). As the fungi grew, an obvious biofilm formed at the fuel:water interface. This growth could provide a physical niche in that the fungi could use the available oxygen in the fuel for metabolism while excluding other microorganisms from that space and define which organisms can grow in fuel storage tanks. Biodiesel and biodiesel blends (such as B20) contain more dissolved oxygen than ultra-low sulfur diesel (41). Aerobic metabolism at the fuel:water interface would deplete the oxygen present in the immediate environment, including the aqueous phase. Dissolved oxygen in the much larger volume of the fuel would be limited by diffusion into the aqueous phase, which would more than likely remain anoxic when contaminated with metabolizing microorganisms. The resulting oxygen concentration gradient would be most prominent at the interface of fuel and water where the fungi we isolated are actively growing within a fuel tank, generating an oxygenic corrosion cell and leading to exacerbated, aggressive pitting-type corrosion (40, 42).

Although correlative links between microbial degradation of both fossil and bio-based fuels and corrosion are well established, the direct connection between fungal taxa and rates or types of corrosion is less developed (43–45). The controlled experiments in this study link active fungal metabolism to the aerobic degradation of B20, the production of biofilms at the fuel/water interface, and pitting corrosion. Future research will address how potential metabolic interactions between *Byssochlamys* and the bacterial populations might affect the rates of fuel degradation and corrosion. The FAME components of biodiesel are more amenable to biodegradation than petroleum-based hydrocarbons (Figure 8), suggesting persistent challenges in the long-term storage of biodiesel (46, 47). While we tested a 20% biodiesel blend, biodiesel is also present in ULSD in the U.S. at concentrations up to 5%; added to compensate for the loss of lubricity in diesel fuel from the removal of organosulfur compounds. Blends in the EU contain up to 7% biodiesel. Therefore, even in fuels denoted as “neat” or ULSD the degradation of FAME and hydrocarbons represents an increased corrosion risk within storage infrastructure. These findings have identified the challenges associated with the incorporation of FAME in diesel fuels and will inform best management practices to allow the continued use of renewable fuels, while also reducing the risk of microbiologically influenced corrosion to global energy infrastructure.

## MATERIALS AND METHODS

### Isolation, identification, and growth of fungal isolates

*Byssochlamys* sp. strain AF004 and *Wickerhamomyces* sp. strain SE3 were isolated from contaminated B20 biodiesel storage tanks at an Air Force Base in the southeast United States (32). To isolate the fungi, 1 L of B20 fuel was passed through a Steritop™bottle top filter unit with a 40 cm^2^ filter area and 0.22 µm nominal pore size (Millipore Sigma). This filter was cut out with a sterile scalpel, transferred to a 1.5 mL Eppendorf tube containing 500 µL of phosphate buffered saline (pH 7.4), and the biomass was resuspended by vortexing at maximum speed for 1 min. The biomass suspension was diluted in phosphate buffered saline (pH 7.4), spread onto Hestrin Schramm (HS) agar medium (per L: 20g glucose, 5g yeast extract, 5g peptone, 2.7g Na_2_HPO_4_, 1.15g citric acid, 7.5g Agar; pH adjusted to 6.0 with diluted HCl or NaOH)(48) and incubated at 25°C for 7 days.

Fungal growth was transferred for isolation repeatedly until pure cultures (repeatedly singular morphologies) were obtained. The fungal isolates were identified by extracting genomic DNA with the *Quick*-DNA Fungal/Bacterial Miniprep Kit (Zymo Research; Irvine (Fig, CA), amplifying and sequencing the internal transcribed spacer (ITS) region. Specifically, the ITS region for each isolate was amplified with the primers ITS1-F (5’CTTGGTCATTTAGAGGAAGTAA-3’) and ITS4 (5’-TCCTCCGCTTATTGATATGC-3’) (49, 50)in a PCR using 5 PRIME HotMasterMix (Quanta Biosciences, Beverly, MA, United States). The amplified DNA was purified using ExoSap-IT™(Thermo Fisher Scientific) according to the manufacturer’s instructions. Once purified, the amplified DNA was submitted for BigDye^®^ Direct Cycle sequencing with each amplification primer on the 3130xl Genetic Analyzer (Biology Core Molecular Laboratory, University of Oklahoma). The resulting sequences for each amplicon were trimmed for quality to only include nucleotides with a Q-score ≥ 30 and merged using CAP3 (51). The trimmed and merged sequences were then submitted to NCBI to identify their closest neighbor using MegaBLAST. The MEGA X software package (52) was used to build a maximum likelihood tree (Tamura-Nei model, bootstrapped with 500 samplings) containing sequences from the isolates and closest neighbors (Fig. 1).

A spore suspension of the filamentous fungus *Byssochlamys* sp. was used as the normalized inoculum in the experiments described below. To prepare the spore suspension, 4 mL of PBS was added to the surface of HS agar containing hyphal growth of *Byssochlamys* sp., and an inoculating loop was used to scrape off the fungal growth. The PBS solution was then collected from the plate and filtered through a 10 µm pore size nitrocellulose filter, to separate spores from hyphal biomass. The spores were then centrifuged at 10,000 x RCF for 1 minute. The supernatant was decanted, and sterilized PBS was added back to the spore pellet and vortexed to resuspend the spores in solution. This was repeated for a total of three washes. Spore concentrations were determined using a Petroff-Hausser counting chamber then diluted to adjust the inoculum concentration to 1×10^4^ spores/mL.

To produce a suspension of yeast cells, *Wickerhamomyces* sp. was grown in HS broth for 48 hours and was centrifuged at 10,000 x RCF to pellet cell mass. The supernatant was decanted, and sterilized PBS was added to the cell pellet. This pellet was centrifuged at 10,000 x RCF for 1 minute. The supernatant was decanted, and more sterilized PBS was added to the cell pellet and vortexed to resuspend the cells in suspension. This was repeated for a total of three washes. Cell concentrations were determined using a Petroff-Hausser counting chamber and diluted to adjust the inoculum concentration to 1×10^4^ cells/mL.

### Quantification and Characterization of B20 Biodiesel Fuel Biodegradation

Fungal biodegradation of B20 was evaluated by direct measure of fungal growth and consumption of the fuel compounds. Fungal growth and ability to degrade B20 biodiesel was measured every 7 days for 21 days. All biodegradation experiments were incubated aerobically at 25 °C and non-shaking (i.e. static) conditions. Test tubes (16 x 150 mm) were filled with 5 mL filter sterilized B20 as the sole carbon and energy source and ASW liquid medium in a 1:10 fuel to ASW ratio. *Wickerhamomyces* and *Byssochlamys* were initially grown on HS broth and spun down at 10,000 x RCF to pellet cell mass. This pellet was resuspended in PBS by vortex, centrifuged and resuspended a total of 3 times to remove any potential nutrient carry over before inoculation. Finally, the cell/spore density was determined using a hemocytometer, allowing the B20 and ASW to be inoculated to a final concentration of 1×10^4^ cells/mL or spores/mL respectfully. The fuel phase of these cultures was separated using a 10 mL separatory funnel and diluted 1:10 in hexane ≥97.0% (GC) and analyzed by Gas Chromatography/Mass Spectrometry (GC/MS). The chemical composition of the B20 biodiesel samples prior to degradation was determined by GC/MS using a Shimadzu QP 2010 SE (Shimadzu Corporation, USA). Each sample was diluted 1:10 with hexane prior to injection. A volume of 1 µL was injected via autosampler with a split ratio of 1:10 for a final dilution of 1:100. Injection started at 300 °C, the oven was at 40°C with a 1.5 min hold, which increased to 320°C at a rate of 10°C min^-1^. Chemical components were separated with a Restek Column Rxi 5Sil with dimensions: 30 m, 0.25 mm ID, 0.25 µm. High purity helium was used as a carrier gas at a linear velocity of 36.8 cm s^-1^. Mass spectra were analyzed in scan mode with the following parameters: interface at 320 °C, ion source 200 °C, solvent cut of 2.75 min, event time of 0.25 sec and scan speed of 2000. Each Total Ion Chromatogram (TIC) was processed using the software LabSolutions version 4.20 (Shimadzu Corporation, USA). Peaks were identified using the mass spectra library NIST version 14 and verified and quantified using reference standards for FAME (Supelco® 37 Component FAME Mix, Sigma Aldrich, USA) and saturated alkanes (C7-C40 Saturated Alkanes Standard, Sigma Aldrich, USA). Major alkane and FAME peaks were identified by the NIST library replicates and underwent destructive sampling of quintuplicates at each time point. The degradation of fuel compounds was measured by determining the amount (%) of the remaining fuel components relative to a non-exposed control. For all experiments, negative controls (un-inoculated) were included to evaluate contamination risks and assess abiotic degradation. Un-amended controls (inoculated but with no B20) were also included to evaluate nutrient carryover from the initial inocula.

### Quantification and Characterization of Microbiologically Influenced Corrosion in Bioreactors

Three CDC biofilm bioreactors^**®**^ (BioSurface Technologies Corp.) were filled with 3 mL of B20 biodiesel and 297 mL of Artificial Sump Water (ASW, per L: 0.015g NaCl, 0.035g NaF, 0.02g CaCl_2_, 0.018g KNO_3_, 0.01g Na_2_SO_4_, 0.015g (NH_4_)_2_SO_4_, and 0.017g K_2_HPO_4_) (1:100 ratio of B20:ASW) and inoculated to a final concentration of 1×10^4^ *Byssochlamys* spores/mL or *Wickerhamomyces* cells/mL. Grade 1018 carbon steel circular disk coupons (1.27 cm diameter x 3.82 mm thick; BioSurface Technologies Corp., Bozeman, MT) were washed in acetone to remove any machine oil, weighed to obtain initial mass, and sterilized prior to use by exposing them to UV light for 15 minutes on each side. In an AirClean 600 PCR Workstation (AirClean Systems Inc., Creedmoor, NC), three carbon steel disks were inserted into each of three polypropylene coupon holders. Three of the coupon holders were inserted into each of the three bioreactors. The bioreactors were sampled weekly for three weeks by removing one of the polypropylene coupon holders and replacing it with a sterilized bioreactor blank coupon holder, allowing for three technical and three biological replicates. The coupons collected from each reactor were cleaned and weighed to determine mass loss, which was used to calculate corrosion rates (described below).

The surface topology of each coupon was also determined to quantify the area and depth of any corrosion. The pH of the aqueous phase was determined at each sampling time, as was the viability of the isolates or sterility of controls. Viability measurements of the planktonic microorganisms were made by sampling medium from the bioreactors and conducting MPN/mL measurements for *Byssochlamys* AF001 and CFU/mL measurements for *Wickerhamomyces* SE3 using liquid or solid HS medium, accordingly. The population density of microorganisms attached to each coupon was determined by removing biomass with a sterile nylon swab and transferring it to sterile PBS. The biomass was suspended by vortexing and used to determine MPN or CFU/mL as described above. After removing the biomass with a swab, each coupon was cleaned using ASTM method G01-03 C3.5 and immediately weighed to determine mass loss. After being weighed, the coupons were stored in an anaerobic chamber to prevent further abiotic oxidation until surface analysis could be conducted. The technical replicates were averaged and used for data analysis.

The surface of each coupon was analyzed using white light profilometry (Nanovea PS50; Nanovea, Inc.; Irvine, CA). To ensure that the same area was being scanned on all coupons, a clip was used to align the coupons in the same location on the profilometer’s stage (Supplemental information contains 3D files that can be 3D printed and examined).

A 6 x 6 mm area was scanned on the coupon and visualized using Mountains Software version 6.3 (Digital Surf). Intensity and height maps were combined in the software and leveled using a least squares plane method by subtraction. Non-measured points were filled in using calculations of points from the nearest neighbors. Pits were defined as any points that were greater than 20 µm below the mean surface average.

Maximum pit depth and total pitted area were calculated from the surface analyses.

### Localizing Corrosion in Fuel, Interphase, and Aqueous Phases

Static corrosion experiments were set up with the intent of determining where corrosion was greatest, in the fuel, interphase, or aqueous phase of water-containing fuel systems. The static incubations were made by first filling test tubes (16 x 150 mm) with 3 mL of sterile ASW medium to mimic water present in the underground storage tanks from runoff or condensation within the tank (53). B20 biodiesel was filter sterilized into an autoclaved 1L Schott Bottle using a Steritop™ bottle top filter unit with a 40 cm^2^ filter area and 0.22 µm nominal pore size (Millipore Sigma) and 3 mL of this biodiesel was aseptically added to the tubes, resulting in a 1:1 fuel to ASW ratio. Non-galvanized carbon steel brads (0.5 mm diameter x 19 mm length) were washed in acetone to remove any machine oil, weighed to obtain initial mass, and autoclaved anaerobically in Balch tubes with a N_2_ headspace to prevent abiotic corrosion and ensure sterility prior to the experiment. The brads were inserted into holes drilled into nylon bolts at three levels before the autoclave cycle. The holes were positioned so that the brads were exposed to either the fuel phase, the fuel-water interface, or the aqueous phase (Supplemental Figure 1).

Five replicates of the static corrosion tubes described above were either inoculated with a final concentration of 1×10^4^ *Byssochlamys* sp. spores or *Wickerhamomyces* sp. cells per mL. An additional five tubes remained uninoculated as controls. The tubes were incubated for 21 days at room temperature in the dark. After 21 days, the pH of the aqueous phase was measured using an Oakton pH Spear waterproof Pocket pH Testr™. The viability of the organisms (or sterility of the controls) was assessed by growth on HS agar medium and direct microscopy. Viability of the *Wickerhamomyces* sp. was determined by CFU/mL on HS agar medium. Growth of the *Byssochlamys* sp. was confirmed by observation of hyphal growth under light microscopy by preparing a wet mount and examining at 100x magnification. Corrosion rates were calculated from mass loss using the approach described below (Calculating corrosion rates from mass loss measurements).

Additional static corrosion experiments were conducted with flat coupons that traversed the fuel, interphase, and aqueous phases in order to visualize the surface of the coupon across all phases with scanning electron microscopy and measure the area and depth of corrosion pits using the white light profilometry as described above. The coupons used for this experiment were composed of 1018 carbon steel, 3” inches long, 3/8” inches wide, and 1/16” inches thick, and finished by the manufacturer by sanding using a 120-grit belt (Alabama Specialty Products, Inc.; Munford, AL). Each coupon was placed in separate 250 mL Erlenmeyer flasks containing 25 mL of sterile B20 biodiesel and ASW for a 1:1 ratio. These flasks were either inoculated with 1×10^4^ spores of the filamentous fungus *Byssochlamys* or kept uninoculated as controls, incubated at 25°C for 90 days, and used to quantify and characterize surface corrosion.

The surface of the coupons was analyzed using white light profilometry as described above. To ensure that the same area 20 x 8 mm area was being scanned on all coupons, a clip was used to align the coupons in the same location on the profilometer’s stage (Supplemental Information).

Intensity and height maps were combined using the Mountains Software version 6.3 (Digital Surf; Besançon, France) and leveled using a least squares plane method by subtraction. Within the software, non-measured points were filled in using calculations of points from the nearest neighbors. Pits were defined as any points that were less than 20 µm below the mean surface average. Maximum pit depth and total pitted area were calculated from the surface analyses.

### Calculation of corrosion rates from mass loss measurements

Following incubation, carbon steel brads and coupons were cleaned using ASTM method G01-03 C3.5 (54). The final mass was measured and used to determine the rates of corrosion in milli-inches per year (MPY) using the following equation:

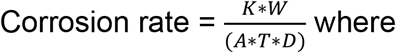

K = Mils per year (MPY) rate

T = Time of exposure in hours

A = Area in cm^2^

W = Mass loss in grams (Initial coupon mass – final coupon mass)

D = Density in g/cm^3^

### Statistical analysis and data visualization

Statistical analysis and figure generation was carried out in R version 3.3.3 and GraphPad Prism 8.3.0. Significant differences were calculated using two-way ANOVA with a Tukey’s HSD to determine significant differences in corrosion rates, maximum pit depths, total pitted areas, and pH between the isolates and uninoculated controls.

## ACKNOWLEDGMENTS

We acknowledge the men and women of the US Air Force and Civilian Personnel at US Air Force bases; their cooperation and assistance were critical to this research. Additionally, we would like to thank Emily Junkins for insightful comments during the development of this manuscript. This work was supported by the Air Force Research Laboratory Biological Materials and Processing Research Team, Materials and Manufacturing Directorate and the U.S. Department of Defense Office of Corrosion Policy & Oversight Technical Corrosion Collaboration (Grant # FA7000-15-2-0001).

## 3D files used to scan witness coupons at precise locations on using the Nanovea PS 50

https://github.com/Jfloydo/3D-Printed-White-Light-Profilometer-Stage-Cover-for-Nanovea-PS50/tree/main

## WhiteLightStage_CorrectedBase_BioreactorCouponsForCircularcouponswithDia mter0.5inches.glb

Supplementary 3D file for printing a cover over the Nanovea PS50 that supports the placement of circular surface coupons with diameters of 12.7mm/0.5in.

## WhiteLightStage_CorrectedBase_3inx0.5in.glb

Supplementary 3D file for printing a cover over the Nanovea PS50 that supports the placement of rectangular surface coupons with lengths of 76.2mm/3in and widths of 12.7mm/0.5in.

